# The diversity, evolution and ecology of *Salmonella* in venomous snakes

**DOI:** 10.1101/526442

**Authors:** Caisey V. Pulford, Nicolas Wenner, Martha L. Redway, Ella V. Rodwell, Hermione J. Webster, Roberta Escudero, Carsten Kröger, Rocío Canals, Will Rowe, Javier Lopez, Neil Hall, Paul D Rowley, Dorina Timofte, Robert A. Harrison, Kate S. Baker, Jay C. D. Hinton

## Abstract

**Background:** Reptile-associated *Salmonella* are a major, but often neglected cause of both gastrointestinal and bloodstream infection globally. The diversity of *Salmonella enterica* has not yet been determined in venomous snakes, however other cold-blooded animals have been reported to carry a broad range of *Salmonella* bacteria. We investigated the prevalence and assortment of *Salmonella* in a collection of venomous snakes in comparison with non-venomous reptiles.

**Methodology/Principle Findings:** We used a combination of selective enrichment techniques and whole-genome sequencing. We established a unique dataset of reptilian isolates to study *Salmonella enterica* species-level evolution and ecology and investigated differences between phylogenetic groups. We observed that 91% of venomous snakes carried *Salmonella*, and found substantial diversity between the serovars (n=58) carried by reptiles. The *Salmonella* serovars belonged to four of the six *Salmonella enterica* subspecies: *diarizonae, enterica*, *houtanae* and *salamae*. Subspecies *enterica* isolates were distributed among two distinct phylogenetic clusters, previously described as clade A (52%) and clade B (48%). We identified metabolic differences between *S. diarizonae, S. enterica* clade A and clade B involving growth on lactose, tartaric acid, dulcitol, myo-inositol and allantoin.

**Significance:** We present the first whole genome-based comparative study of the *Salmonella* bacteria that colonise venomous and non-venomous reptiles and shed new light on *Salmonella* evolution. The findings raise the possibility that venomous snakes are a reservoir for human Salmonellosis in Africa. The proximity of venomous snakes to human dwellings in rural Africa may result in contaminated faecal matter being shed on surfaces and in water sources used for human homes and to irrigate salad crops. Because most of the venomous snakes had been captured in Africa, we conclude that the high level of *Salmonella* diversity reflects the African environmental niches where the snakes have inhabited.

**Author Summary:** *Salmonella enterica* is a remarkable bacterial species that causes Neglected Tropical Diseases globally. The burden of disease is greatest in some of the most poverty-afflicted regions of Africa, where salmonellosis frequently causes bloodstream infection with fatal consequences. The bacteria have the ability to colonise the gastrointestinal tract of a wide range of animals including reptiles. Direct or indirect contact between reptiles and humans can cause Salmonellosis. In this study, we determined the prevalence and diversity of *Salmonella* in a collection of African venomous snakes for the first time. Using the power of genomics, we showed that the majority of venomous snakes (91%) carry *Salmonella*, two thirds of which belonged to a subspecies of *S. enterica* called *enterica*, which is associated with most cases of human salmonellosis. Within the *S. enterica* subspecies we identified two evolutionary groups which display distinct growth patterns on infection relevant carbon sources. Our findings could have particular significance in Africa where venomous snakes wander freely around human dwellings and potentially shed contaminated faecal matter in water sources and on surfaces in rural homes.

## Introduction

*Salmonella* is a clinically relevant bacterial pathogen of great public health significance [1,2]. The *Salmonella* genus contains two species; *S. bongori* and *S. enterica. S. enterica* is further divided into six subspecies; *enterica* (I), *salamae* (II), *arizonae* (IIIa), *diarizonae* (IIIb), *houtanae* (IV) and *indicia* (VI) [3]. The subspecies are classified into approximately 2 600 serovars which are ecologically, phenotypically and genetically diverse [4]. Serovars which belong to *S. enterica* subspecies *enterica* cluster phylogenetically into two predominant clades (A and B) [5–7]. Henceforth *Salmonella* will refer to the *S. enterica* species only, unless stated otherwise.

At the genus level, *Salmonella* has a broad host-range whilst individual serovars differ in host-specificity [8]. The majority of *Salmonella* infections in humans (99%) are caused by a small number of serovars belonging to the *S. enterica* subspecies [9]. Serovars which belong to *non-enterica* subspecies are associated with carriage in cold-blooded animals such as non-venomous reptiles and amphibians, but are rarely found in humans [8,10–12]. Carriage rates of *non-enterica* serovars in reptiles can be high, for example, one study focused on snakes in a pet shop found that 81% of animals were carrying *S. diarizonae* [12]. Further examples of previous studies demonstrating the diverse range of *Salmonella* serovars that colonise various reptilian species in different countries are shown in Table 1.

**Table 1.** The distribution of *Salmonella* subspecies in non-venomous reptiles

| Citation | Country of study | Reptile information | Distribution |
| --- | --- | --- | --- |
| Piasecki *et al.*, 2014 [13] | Poland | Snakes, Lizards, Chelonians | 59% *enterica*<br>16% *salamae* or *houtanae*<br>3.5% *diarizonae* or *indica* |
| Lukac *et al.*, 2015 [14] | Croatia | Snakes, Lizards, Chelonians | 34.6% *enterica*<br>23.1% *houtanae*<br>23.1% *arizonae*<br>15.4% *diarizonae*<br>2.8% *salamae* |
| Nakadai *et al.*, 2005 [15] | Japan | Snakes, Lizards, Turtles | 62.5% *enterica*<br>37.5% *salamae*, *arizonae*, *diarizonae* or *houtanae* |
| Geue and Löschener, 2002 [16] | Germany and Austria | Reptiles | 52.8% *enterica*<br>34.5% *diarizonae*<br>3.4% *salamae*<br>6.9% *arizonae*<br>2.3% *houtanae* |
| Sá and Solari, 2001 [17] | Brazil | Brazilian and Imported Pet Snakes, Lizards and Chelonians | 44.7% *enterica*<br>10.5% *salamae*<br>5.2% *arizonae*<br>21% *diarizonae*<br>18.5% *houtanae* |
| Perderson *et al.*, 2009 [18] | Denmark | Captive reptiles | 65% *enterica*<br>12% *diarizonae*<br>11% *houtanae*<br>6% *salamae*<br>6% *arizonae* |
| Schröter *et al.*, 2004 [12] | Germany | Captive reptiles | 100% *diarizonae* |

Reptiles represent a significant reservoir for serovars of *Salmonella* associated with human disease, as over 60% of reptiles have been reported to carry serovars falling within the *S. enterica* subspecies [18]. About 6% of human salmonellosis cases are contracted from reptiles in the USA [19], and in South West England, 27.4% of *Salmonella* cases in children under five years old were linked to reptile exposure [20]. The latter study demonstrated that reptile-derived salmonellosis was more likely to cause bloodstream infection in humans than non-reptile-derived *Salmonella* [20]. Reptile-associated *Salmonella* is therefore considered to be a global threat to public health [21].

The majority of reptile-associated salmonellosis cases reported in humans are caused by *Salmonella* from non-venomous reptiles [21], most likely because these are more frequently kept as pets. Therefore, non-venomous reptiles have been the focus of numerous studies while the prevalence and diversity of *Salmonella* in venomous snakes remained unknown. The evolution of venom function is a key factor that drives ecological diversification in snakes [22], with venomous snakes differing from non-venomous reptiles in a number of important aspects including the way that animals interact with the environment, food sources, habitat and humans which may have concomitant effects upon *Salmonella* carriage.

Research to improve snakebite treatment at the Liverpool School of Tropical Medicine (LSTM) has resulted in the creation of the UK’s most extensive collection of venomous snakes (195). The LSTM herpetarium houses venomous snakes from a diverse range of species and geographical origins representing an ideal source of samples to assess *Salmonella* in this under-studied ecological niche. We collected *Salmonella* isolates from 97 venomous snakes housed at LSTM, and made genome-level comparisons with *Salmonella* collected from 28 non-venomous reptiles from the University of Liverpool veterinary diagnostic laboratory.

The aims of this study were three-fold. Firstly, to determine the period prevalence of *Salmonella* in a collection of captive venomous snakes and investigate whether this group of reptiles are potential reservoirs for human salmonellosis in Africa. Secondly, to assess the serological and phylogenetic diversity of *Salmonella* amongst venomous snakes compared with non-venomous reptiles. Thirdly, to utilise the diversity of *Salmonella* carried in both venomous snakes and non-venomous reptiles to assess clade-specific differences thought to reflect adaptation to survival in the environment or different hosts. Here, we present the first whole genome-based comparative study of the *Salmonella* bacteria that colonise venomous and non-venomous reptiles.

## Methods

### Source of *Salmonella* isolates

The *Salmonella* isolates were derived from faecal samples from two collections of reptiles. One hundred and six faecal samples were collected from venomous snakes at LSTM from May 2015 to January 2017, with an emphasis on snakes originating from Africa (S1 Table), and investigated for the presence of *Salmonella*. All venomous snakes where housed in individual enclosures and fed with frozen mice. Sixty-nine of the samples (71%) were sourced from wild-caught snakes originating from: Togo, Nigeria, Cameroon, Egypt, Tanzania, Kenya, South Africa, and Uganda. A further 28 *Salmonella* isolates (29%) came from venomous snakes bred in captivity. The LSTM herpetarium is a UK Home Office licensed and inspected animal holding facility. A second collection of 28 *Salmonella* isolates from non-venomous reptiles were sourced from the veterinary diagnostics laboratory based at the University of Liverpool’s Leahurst campus (reptilian species described in S1 Table). Leahurst isolate 11L-2493 was isolated from a beaded lizard which has the ability to produce venom. These isolates were collected from June 2011 to July 2016 from specimens submitted as part of *Salmonella* surveillance for import/export, but also clinical faecal samples and tissues from post mortem investigations. The provenance of the non-venomous reptile isolates is described in S1 Table. The majority of the non-venomous reptiles were sourced from a zoological collection, however two animals were privately owned and three were sourced from the RSPCA. The LSTM isolates are henceforth referred to as venomous snake isolates and the Leahurst isolates are referred to as non-venomous reptile isolates.

### Isolation of *Salmonella*

All media were prepared and used in accordance with the manufacturers guidelines unless otherwise stated. *Salmonella* was isolated using a modified version of the protocol described in the national Standard Operating Procedure for detection of *Salmonella* issued by Public Health England [23].

Faecal droppings were collected from reptiles and stored in 15 mL plastic centrifuge tubes at 4°C. Two different methods were used for the enrichment of *Salmonella* from faecal samples due to reagent availability at the time of isolation. S1 Table provides information on isolate specific methods. In enrichment method 1, faecal samples were added to 10 ml of buffered peptone water (Fluka Analytical, UK, 08105-500G-F) and incubated overnight at 37°C with shaking at 220 rpm. Following overnight incubation, 100 μL of the faeces mixture was added to 10 mL of Selenite Broth (19 g/L selenite broth base, Merck, UK, 70153-500G and 4 g/L sodium hydrogen selenite, Merck 1.06340-50G) and incubated overnight at 37°C with shaking at 220 rpm. In enrichment method 2, faecal samples were added to 10 ml of Buffered Peptone Water (Fluka Analytical, 08105-500G-F) supplemented with 10 μg/ml Novobiocin (Merck, N1628), and incubated overnight at 37°C with shaking at 220 rpm. Following overnight incubation, 100 μL of the faeces mixture was added to 10 ml Rappaport-Vassilliadis Medium (Lab M, UK, LAB086) and incubated for 24 hours at 42°C with shaking at 220 rpm.

Following enrichment, 10 μL of overnight broth was spread onto Xylose Lysine Deoxycholate (XLD) (Oxoid, UK, CM0469) agar plates which were incubated overnight at 37°C. Putative *Salmonella* colonies were selected by black appearance on XLD plates and confirmed by pink and white colony formation on Brilliant Green Agar (Merck, 70134-500G) supplemented with 0.35 g/L mandelic acid (Merck, M2101) and 1 g/L sodium sulfacetamide (Merck, S8647).

To identify *S. enterica* species, colony PCR of the *Salmonella* specific *ttr* locus, which is required for tetrathionate respiration [24], was performed. PCR reagents included MyTaq Red Mix 1x (Bioline, UK, BI0-25043), *ttr-4* reverse primer (5’-AGCTCAGACCAAAAGTGACCATC-3’) and *ttr-6* forward primer (5’-CTCACCAGGAGATTACAACATGG-3’) on colonies suspected to be *Salmonella*. PCR reaction conditions were as follows: 95°C 2 min, 35 x (95°C 15 s, 60°C 30 s, 72°C 10 s), 72°C 5 min. PCR products were visualised using agarose gel (3.5%) (Bioline, BIO-41025) electrophoresis in TAE buffer. Midori Green DNA stain (3 μL/100 mL) (Nippon Genetics, Germany, MG 04) was used to visualise DNA bands under UV light. Throughout the isolation procedure, *S. enterica* serovar Typhimurium (S. Typhimurium) strain LT2 [25,26] was used as a positive control, and *Escherichia coli* MG1655 [27] was used as a negative control (S1 Table).

### Whole-genome sequencing of *Salmonella* from venomous and non-venomous reptiles

All non-venomous reptile isolates and 87 of 97 venomous snake isolates were sent for whole genome sequencing. LSS-88 to LSS-97 were not sequenced as they were collected after the submission deadline to the sequencing centre. Isolates were sent to either MicrobesNG, UK or the Earlham Institute, UK for whole-genome sequencing on the Illumina HiSeq platform (Illumina, California, USA). Isolates which were sequenced by MicrobesNG were prepared for sequencing in accordance with the company’s preparation protocol. Isolates which were sequenced by the Earlham Institute were prepared by inoculating a single colony of *Salmonella* into a FluidX^®^ 2D Sequencing Tube (FluidX Ltd, UK) containing 100 μL of Lysogeny Broth (LB, Lennox) and incubating overnight at 37°C, with shaking at 220 rpm. LB was made using 10 g/L bacto tryptone (BD Biosciences, UK, 211705), 5 g/L bacto yeast extract (BD, 212750) and 5 g/L sodium chloride (Merck, S3014-1kg). Following overnight growth, the FluidX^®^ 2D Tubes were placed in a 95°C oven for 20 minutes to heat-kill the isolates.

DNA extractions and Illumina library preparations were conducted using automated robots at MicrobesNG or the Earlham Institute. At the Earlham institute, the Illumina Nextera XT DNA Library Prep Kit (Illumina, FC-131-1096) was used for library preparation. High throughput sequencing was performed using an Illumina HiSeq 4 000 sequencing machine to achieve 150 bp paired-end reads. Sequencing was multiplexed to 768 unique barcode combinations per sequencing lane. The insert size was determined to be approximately 180 bp and the median depth of coverage was 30x.

At MicrobesNG, genomic DNA libraries were prepared using the Nextera XT Library Prep Kit (Illumina, FC-131-1096) with two nanograms of DNA used as input and double the elongation time of that described by the manufacturer. Libraries were sequenced on the Illumina HiSeq 2 500 using a 250 bp protocol.

### Obtaining contextual reference sequences

Reads were uploaded to Enterobase for serovar prediction using the *Salmonella in silico* typing resource (SISTR) [28,29]. Enterobase was also used to assign a Multi Locus Sequence Type (MLST) to each isolate, based on sequence conservation of seven housekeeping genes [29]. Where available, reference isolates representing previously sequenced *Salmonella* isolates for all subspecies and serovars identified were included in the analysis. Reference sequence assemblies were downloaded from the National Center for Biotechnology Information (NCBI). Accession numbers are available in S2 Table.

### Quality control checks

Fastqc v0.11.5 (www.bioinformatics.babraham.ac.uk/projects/fastqc/) and multiqc v1.0 (http://multiqc.info) were used to assess read quality. Kraken v0.10.5-beta [30] was run to ensure reads were free from contamination using the MiniKraken 8gb database and a *Salmonella* abundance cut-off of 70%. Trimmomatic v0.36 [31] was then used on the paired end reads to trim low-quality regions using a sliding widow of 4:200. ILLUMINACLIP was used to remove adapter sequences.

### Phylogenetics-core genome alignment tree

Genomes were assembled using SPADES v3.90 [32]. QUAST v4.6.3 [33] was used to assess the quality of assemblies, the results of which can be found in S3 Table. Assemblies which comprised of greater than 500 contiguous sequences were deemed too fragmented for downstream analysis. All assemblies which passed QC were annotated using Prokka v1.12 [34]. Roary v3.11.0 [35] was used to generate a core genome alignment. SNP-sites v2.3.3 [36] was used to extract SNPs. A maximum likelihood tree was built from the core genome SNP alignment of all isolates using RAxML-NG v0.4.1 BETA [37] with the general time reversible GTR model and gamma distribution for site specific variation and 100 bootstrap replicates to assess support. The tree was rooted using the *Salmonella* species *S. bongori*. Interactive Tree Of Life v4.2 [38] was used for tree visualisation. We confirmed that there was no bias in phylogenetic signal between the two different sequencing platforms used by assessing clustering patterns within the phylogenies. S1 Table contains details of the sequencing facility. Monophyletic clustering of isolates was used to assign subspecies to newly sequenced *Salmonella* from venomous and non-venomous reptilian hosts. The level of association between venom status and phylogenetic clade was determined using odds ratios and X^2^ statistics using the OpenEpi website (http://www.openepi.com).

### Identification of clade-specific genomic regions

Genes involved in the utilisation of each carbon source were identified using KEGG [39]. Sequences were downloaded using the online tool SalComMac [40], which allows the download of fasta sequences of the genes in *S*. Typhimurium strain 4/74. In the case of *lac* genes the sequences were taken from the *E. coli* reference sequence MG1655. The sequences can be found in S1 Text.

The software tool MEGABLAST v2.2.17 [41] was used to perform a BLAST search of genes in the reptile-derived genomes against a custom-made database of genes diagnostic of *Salmonella* Pathogenicity Islands and genes involved in carbon utilisation. To confirm all MEGABLAST results, the short reads were mapped against each gene using BWA v0.7.10 [42] and SAMtools v0.1.19 [43]. The resulting bam files were manually assessed for gene presence and absence using Integrative Genomics Viewer v2.4.15 [44]. The results were plotted against the maximum likelihood phylogeny using Interactive Tree Of Life v4.2 [38].

### Carbon source utilisation

Differential carbon source utilisation of 39 reptile-derived *Salmonella* isolates from *S. diarizonae, S. enterica* clade A and *S. enterica* clade B was assessed. Filter-sterilised carbon sugar solutions were added into M9 (Merck, M6030-1kg) agar at concentrations which are described in S4 Table. Isolated colonies were transferred from LB agar plates onto M9 carbon source plates using a sterile 48-pronged replica plate stamp and incubated at 37°C firstly under aerobic conditions. If no growth was seen under aerobic conditions for a carbon source, then the procedure was repeated under anaerobic conditions (approx. 0.35% oxygen) with 20 mM Trimethylamine N-oxide dehydrate (TMAO) (Merck, 92277) as a terminal electron acceptor. Anaerobic conditions were achieved by incubating plates in an anaerobic jar with 3x AnaeroGen 2.5 L sachets (Thermo Scientific, UK, AN0025A) to generate anaerobic gas. Oxygen levels were measured using SP-PSt7-NAU Sensor Spots and the Microx 4 oxygen detection system (PreSens, Regensburg, Germany). An LB control plate was used to validate successful bacterial transfer and all experiments were performed in duplicate. *Salmonella* growth was determined at 18, 90 and 162 hours in aerobic growth conditions and at 162 hours in anaerobic growth conditions. A sub-set of growth positive isolates were assessed for single colony formation to validate the results of the replica plating.

### Antimicrobial susceptibility testing of *Salmonella* isolated from venomous and non-venomous reptiles

Antimicrobial susceptibility was determined using a modified version of the European Committee on Antimicrobial Susceptibility Testing (EUCAST) disk diffusion method [45] using Mueller Hinton (Lab M, LAB039) agar plates and a DISKMASTER ^™^ dispenser (Mast Group, UK, MDD64). Inhibition zone diameters were measured and compared to EUCAST zone diameter breakpoints for Enterobacteriaceae [46]. Isolates were first tested with six commonly used antibiotics (Ampicillin 10 μg, Chloramphenicol 30 μg, Nalidixic Acid 30 μg, Tetracycline 30 μg, Ceftriaxone 30 μg, and Trimethoprim/Sulfamethoxazole 25 μg) and were then tested with five additional antibiotics (Meropenem 10 μg, Gentamicin 10 μg, Amoxicillin/Clavulanic Acid 30 μg, Azithromycin 15 μg, and Ciprofloxacin 5 μg) if any resistance was seen to the primary antibiotics (all disks from Mast Group). If resistance was observed phenotypically, then the presence of antimicrobial resistance genes were investigated using the Resfinder software tool [47]. Antimicrobial resistance was defined as resistance to one antimicrobial to which isolates would normally be susceptible [48]. Multidrug resistance was defined as an isolate which showed resistance to three or more antimicrobials to which it would normally be susceptible [48].

### Data availability

Fastq files are available at NCBI Short Read Archive under the accession numbers listed in S1 Table (BioProject Number PRJNA506152, SRA Study Number SRP170008.)

## Results and Discussion

### *Salmonella* prevalence in venomous snakes is similar to that identified in non-venomous reptiles

*Salmonella* carriage is well documented amongst reptiles (Table 1), however, to our knowledge there is no published study that reports on *Salmonella* in venomous snakes. The period prevalence of *Salmonella* was assessed in a collection of 106 venomous snakes housed at the LSTM venom unit between May 2015 and January 2017. A remarkably high proportion (91%; 97/106) of the faecal samples contained *Salmonella* (S1 Table), which should be seen in the context of the significant carriage rate of *Salmonella* by other non-venomous reptiles described in the literature [21]. Our findings pose important public health considerations for individuals who work with venomous snakes, which may previously have been overlooked.

### *Salmonella* diversity in venomous snakes and non-venomous reptiles highlights the possibility of local transmission events and long-term shedding

To assess diversity, 87 venomous snake-derived *Salmonella* isolates and 28 non-venomous reptile-derived *Salmonella* isolates were whole genome sequenced. *In silico* serotyping revealed 58 different *Salmonella* serovars (Fig 1). Multi-locus sequence typing was used to perform sub-serovar genetic characterisation of *Salmonella* [29]. Isolates falling within the same serovar had identical sequence types (S1 Table), reflecting the intra-serovar homogeneity of the *Salmonella* isolated in this study. Taking into account the total number of samples assessed, serovar diversity was greater in non-venomous reptiles (n of serovars = 22, n of samples = 28) compared to venomous snakes (n of serovars = 38, n of samples = 87).

**Fig 1.**
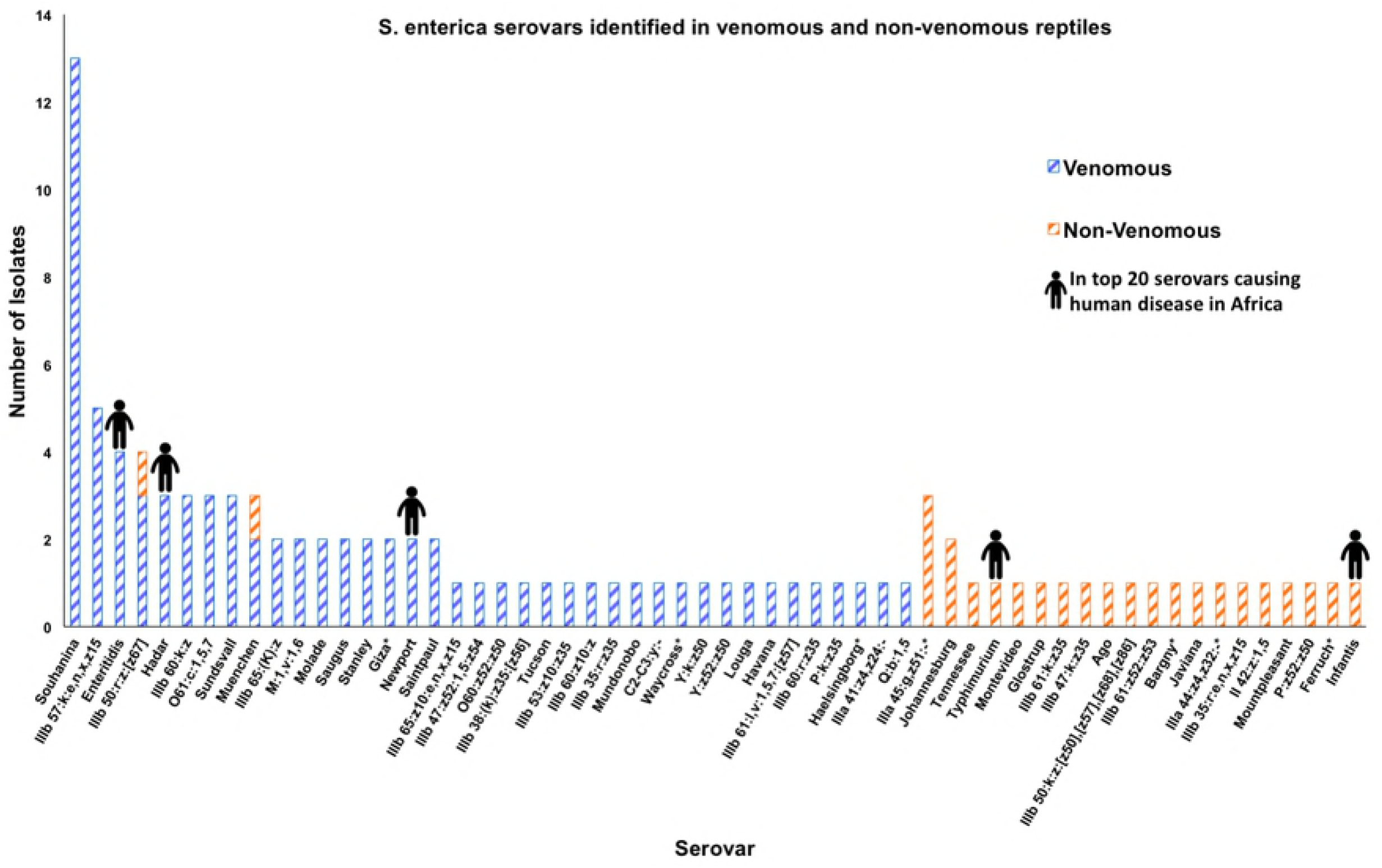
The distribution of the 58 *Salmonella enterica* serovars isolated from venomous and non-venomous reptiles. Each bar represents the total number of isolates which belonged to each serovar. Serovars containing isolates that had multiple serovar designations from SISTR are indicated with asterisks. Human pictographs are displayed on serovars which are amongst the top 20 isolated from humans in Africa. Data is based on the global monitoring of *Salmonella* serovar distribution from the WHO global foodborne infections network data [49].

The most common serovar to be identified in venomous snakes was *S*. Souhanina (n = 15). All *S*. Souhanina isolates cluster locally on the phylogeny (Fig 2) falling within a 5 SNP cluster characteristic of a clonal expansion event [50]. Four of these isolates were found in captive bred reptiles, whilst 11 isolates came from venomous snakes which originated in Cameroon, Uganda, Tanzania, Nigeria, Togo and Egypt. The close phylogenetic relationship between *S*. Souhanina isolates which belong to the same MLST type (ST488) from captive and imported animals with a range of origins suggests that some local *Salmonella* transmission may be occurring. Transmission of *Salmonella* between captured reptiles has been associated with consumption of contaminated food sources such as frozen mice [51,52]. However, a single food source would not explain the high prevalence of venomous snakes which carried distinct and diverse serovars and MLST types.

**Fig 2.**
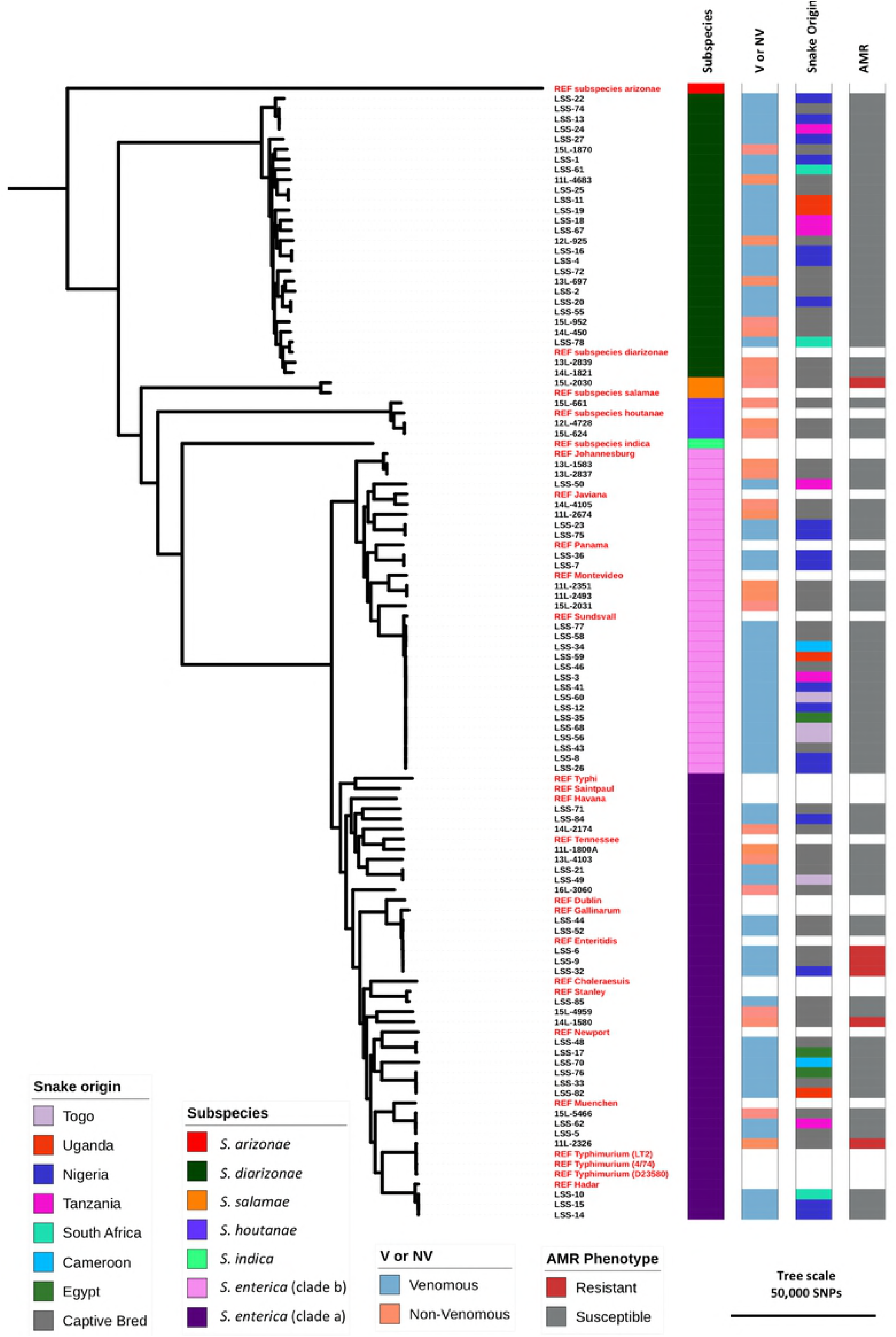
The diversity of *Salmonella* isolated from a collection of venomous snakes and non-venomous reptiles. Core genome maximum likelihood phylogenetic tree. Tree was rooted using *S. bongori* – not shown (S1 Figure). 25 contextual reference genomes representing previously sequenced isolates from each *Salmonella* subgroup are indicated in red. Colour strips showing metadata are as follows; Subspecies – Subspecies of isolate, Origin – The country of origin of the snake from which the isolate was taken, V or NV – Depicts whether the reptile host was venomous (V) or non-venomous (NV), AMR phenotype-Isolates shown in red were resistant to one or more antimicrobial agent. Metadata for the contextual reference genomes appear as white. Tree was visualised using ITOL (https://itol.embl.de).

As long term intermittent shedding of *Salmonella* from reptiles has previously been described [54], we assessed the continuity of *Salmonella* shedding from venomous snakes, by collecting three faecal samples from a Western Green Mamba from Togo over a three-month period between 31^st^ October 2016 and 31^st^ January 2017. All three faecal samples contained *Salmonella* which belonged to sequence type ST488, showing that individual snakes can shed the same sequence type of *Salmonella* over a reasonably long period of time. This continuous shedding of *Salmonella* by a venomous snake demonstrates the risk that reptiles pose to public health. Many of the snakes had been maintained in the LSTM herpetarium for years after collection from the wild. The diverse range of serovars which were isolated suggests that the venomous snakes in this study acquired *Salmonella* in Africa, prior to entering the unit and can shed the bacteria for many years.

### Venomous reptiles carry multidrug resistant *Salmonella* serovars of clinical relevance

Because the majority of venomous snakes examined in this study were of African origin or belonged to a species of snake native to the African continent (Fig 2), we compared the *Salmonella* serovars isolated from all reptiles in this study with those most frequently associated with causing human disease in Africa. The *Salmonella* serovar distribution has been reported by the WHO global foodborne infections network data bank based on data from quality assured laboratories in Cameroon, Senegal and Tunisia [49] (Fig 1). Eleven isolates belonged to serovars commonly pathogenic in humans. We therefore determined the proportion of all venomous snakes and non-venomous reptiles that carried antimicrobial resistant *Salmonella* (Table 2). In *Salmonella* collected from venomous snakes, 4.1% of isolates (4/97) were resistant to at least one antimicrobial and two isolates were multidrug resistant (Table 2). Three resistant isolates from venomous snakes belonged to the serovar Enteritidis and were closely related to the global *S*. Enteritidis epidemic clade which has been shown to be associated with causing human disease in Africa [55]. One of the *S*. Enteritidis isolates (LSS-32) was isolated from a snake which had been imported from Nigeria. These findings raise the possibility that venomous snakes could spread *Salmonella* in African settings where snakes wander freely around the wild and often inhabit rural homes, particularly during the night [56–58]. Such speculation would require in-field epidemiological validation. However, working alongside wild venomous snakes does present logistical challenges.

**Table 2.**
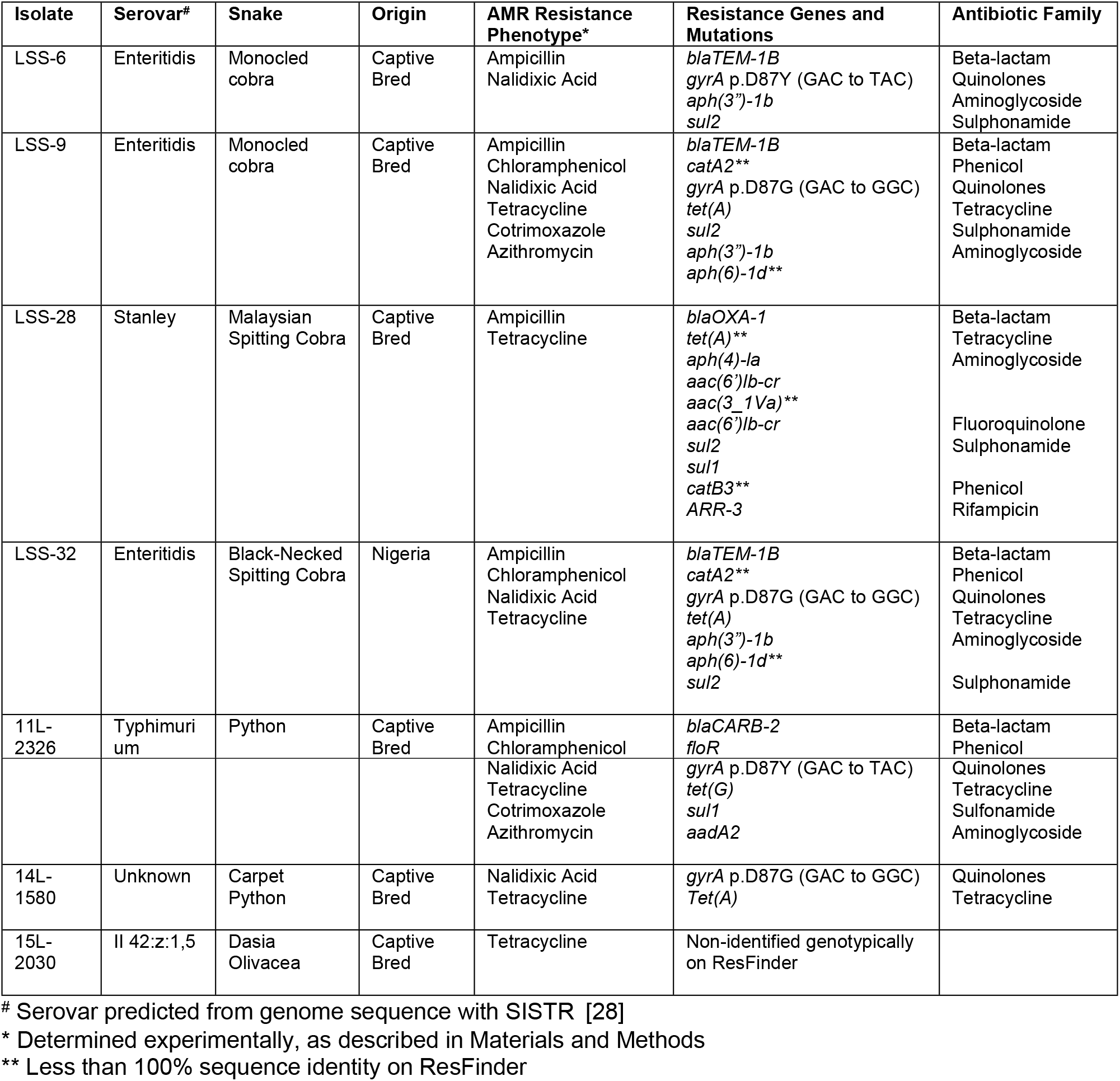
Relating antimicrobial resistance to phenotype and genotype

### Phylogenetic diversity and molecular epidemiology of *Salmonella* in venomous snakes and non-venomous reptiles demonstrated for the first time

By studying *Salmonella* isolated from venomous and non-venomous reptiles we have shown that venomous snakes can shed *Salmonella*. The vast diversity of *Salmonella* has long been acknowledged in the literature [29], however it is rare to see such a broad representation of serovars in a single ecological niche. To study the diversity of reptile-associated *Salmonella* from an evolutionary perspective, we obtained 87 high quality whole genome sequences for the phylogenetic comparison with 24 contextual *Salmonella* genomes (methods). These included 60 isolates from venomous snakes, which were compared with 27 *Salmonella* isolates from non-venomous reptiles following a comprehensive comparative genomic analysis. We identified a total of 405 231 core genome SNPs that differentiated the 87 isolates, and were used to infer a maximum likelihood phylogeny (Fig 2). SNPs are considered to be a valuable marker of genetic diversity [50], and the hundreds of thousands of core-genome SNPs reflect a high level of genetic diversity between the reptile associated *Salmonella* isolates. The collection of reptile-derived *Salmonella* represented most of the known diversity of the *Salmonella* genus [3], spanning four of the six *Salmonella enterica* subspecies: *diarizonae, enterica, houtanae* and *salamae*. Reptile-derived *S. enterica* subspecies *enterica* isolates were approximately equally distributed among into two distinct phylogenetic clusters, known as clade A (58%) and clade B (48%) [5–7] (Fig 2). No significant association was found between venom status and phylogenetic group (OR = 1.1, CI = 0.3-3.0, χ^2^ = 0.02, P = 0.4).

### Genotypic and phenotypic conservation of infection-relevant carbon source utilisation and virulence associated genomic regions sheds new light on *Salmonella* ecology

The unique collection of diverse *Salmonella* isolates was used to determine the phenotypic and genotypic conservation of infection-relevant properties and genomic elements. Whilst the reptile-associated *Salmonella* belonged to five evolutionary groups, the majority of isolates were classified as *S. diarizonae* or *S. enterica*. The clustering of *S. enterica* into two clades (A and B) has previously been inferred phylogenetically based on the alignment of 92 core loci [5,6]. The biological significance of *S. enterica* clade A and clade B has been established as the two clades differ in host specificity, virulence-associated genes and metabolic properties such as carbon utilisation [7]. The genome sequences were used to expand upon pre-existing knowledge and determine phenotypic and genotypic conservation of metabolic and virulence factors across *S. diarizonae* and *S. enterica* (clades A and B).

Although the majority of *Salmonella* serovars of public health significance belong to clade A, certain clade B serovars such as *Salmonella* Panama have been associated with invasive disease [59,60]. Clade B *S. enterica* generally carry a combination of two *Salmonella* genomic islands: *Salmonella* Pathogenicity Island-18 and the cytolethal distending toxin islet. It has been suggested that the two islands are associated with invasive disease, as previously they had only been identified in *S. enterica* serovar Typhi and Paratyphi A, which cause bloodstream infections [1,5]. The combination of *hylE* and *cdtB* genes were present in all *S. diarizonae* and *S. enterica* clade B isolates in this study, but absent from all but one *S. enterica* clade A isolates (14L-2174). We propose that the significant proportion of reptiles which carried *S. enterica* clade B could partially explain the increased likelihood of snake-associated salmonellosis causing invasive disease, compared to non-snake-acquired salmonellosis [20].

The ability of *Salmonella* to grow in a wide range of conditions reflects the adaptation of the bacteria to survival in the environment or in different hosts, as demonstrated by a recent study focused on genome-scale metabolic models for 410 *Salmonella* isolates spanning 64 serovars in 530 different growth conditions [61]. We were interested in examining the conservation of serovar-specific metabolic properties amongst the subspecies and clades of *Salmonella* identified here, and the extent by which genotype reflected growth phenotype. To assess metabolic differences between *S. enterica* clade A, *S. enterica* clade B and *S. diarizonae*, we phenotypically screened 39 reptile isolates for the ability to catabolise a number of infection-relevant carbon sources [55,62–64] (S4 Table). A summary of the results for phenotypic carbon utilisation and the presence of genes associated with the cognate metabolic pathway is shown in Fig 3.

**Fig 3.**
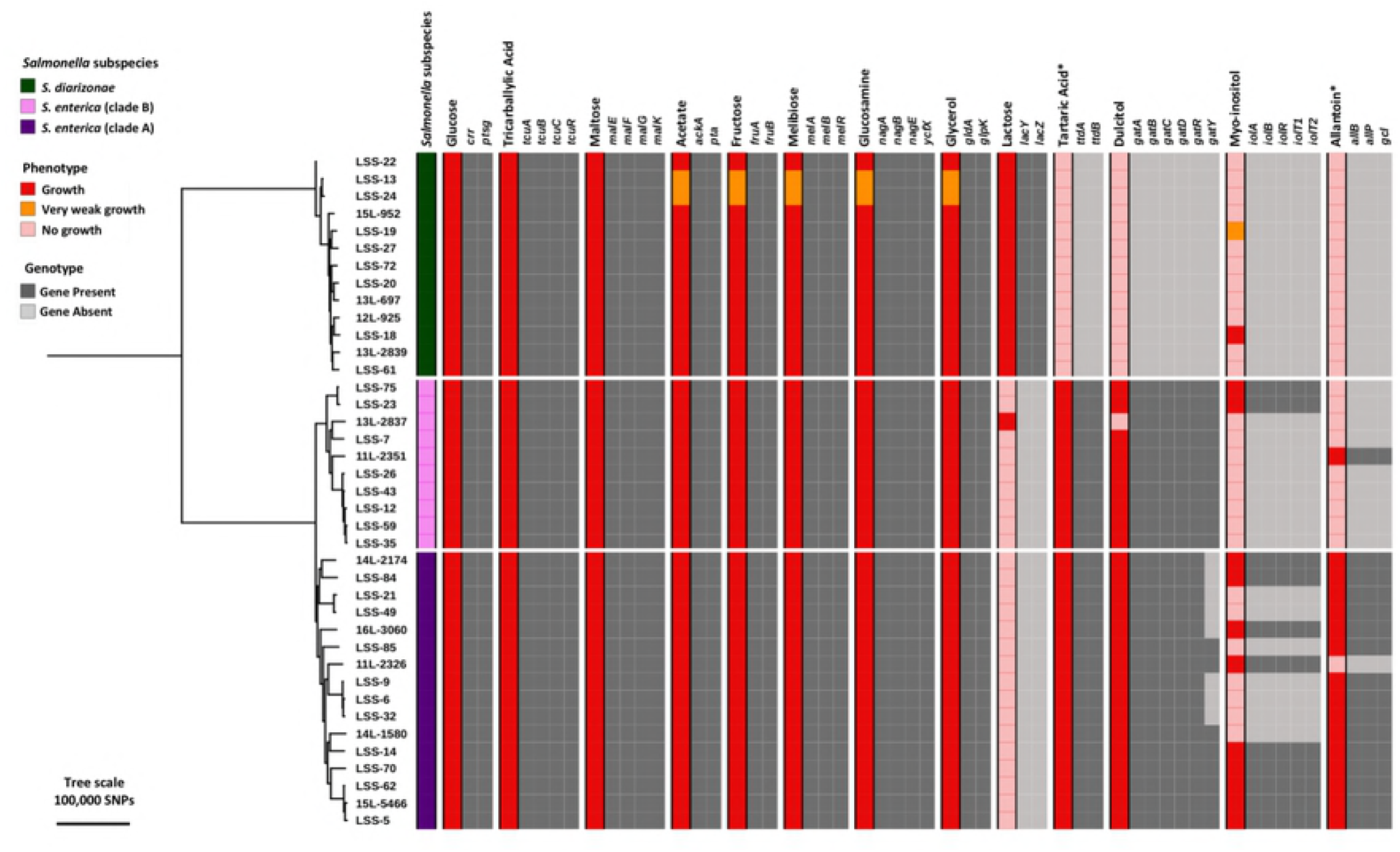
Phylogenetic context of carbon-source utilisation by reptile-derived-Salmonella isolates. Carbon source data were mapped against the core genome phylogenetic tree. The maximum likelihood tree includes all snake-derived *Salmonella* isolates which were assessed for carbon utilisation and had high quality genome sequences. Reference sequences for the majority of carbon genes were taken from *S*. Typhimurium strain 4/74, *lac* gene sequences were taken from *E. coli* MG1655. Carbon sources which required anaerobic conditions are indicated with asterisk.

In general, genotype accurately reflected phenotype in terms of carbon source utilisation; however, this was not always the case (Fig 3). Discrepancies between phenotypic growth and genotype suggests that mechanisms of *Salmonella* metabolism remain to be described. For example, *S. diarizonae* isolate LSS-18 grew well on myo-inositol as a sole carbon source (Fig 3) but showed zero percent homology with any *iol* genes present in the well-characterised *Salmonella* strain 4/74. The ability to utilise lactose was a property of most *S. diarizonae* isolates, consistent with previous reports that 85% of *S. diarizonae* are *Lac*^+^ [65]. It is estimated that less than 1% of all *Salmonella* ferment lactose due to the loss of the *lac* operon from the *S. enterica* subspecies [66]. It was interesting to discover that one non-venomous snake isolate (13L-2837) which belongs to *S. enterica* clade B was capable of utilising lactose as a sole carbon source. Isolate 13L-2337 belongs to the serovar *S*. Johannesburg and to our knowledge this is the first published occurrence of a Lac^+^ *S*. Johannesburg isolate. The 13L-2837 genome had zero percent homology to any *lac* genes that corresponded to reference strain *E. coli* MG1655 (sequence in S1 Text) (results in Fig 3), suggesting an alternative method for lactose utilisation. The 13L-2837 *S*. Johannesburg isolate also lacked the ability to grow on dulcitol, despite possessing all of the relevant *gat* genes, raising the possibility of an inverse relationship between the ability of *Salmonella* to utilise dulcitol and lactose as a sole carbon source.

The majority of *S. enterica* isolates from both clades utilised dulcitol, whereas dulcitol was rarely used as a sole carbon source by *S. diarizonae*. These findings are consistent with those of a study in Australian sleepy lizards, which demonstrated that dulcitol utilisation was observed in almost all *S. enterica* and *S. salamae* isolates but only 10% of *S. diarizonae* isolates [67]. Over 50% of the *S. enterica* clade A isolates lacked the *gatY*gene but grew well on dulcitol as a sole carbon source. The *gatY* gene does not appear to be vital for dulcitol catabolism. Differential repertoires of dulcitol catabolic genes across *Salmonella* serovars are described by Nolle *et al*. (2017) [68], who found that *Salmonella* carries one of two *gat* gene clusters. However, both of these clusters carry the *gatY* gene. Our findings may indicate the possibility that a third *gat* gene cluster, is carried by some *Salmonella* serovars.

In the majority of cases, allantoin was only utilised as a sole carbon source by *S. enterica* clade A isolates, consistent with a previous report that described an association of clade A with the allantoin catabolism island [5]. The ability of *Salmonella* to utilise allantoin is considered to be an adaptation to mammalian and avian hosts, as allantoin is the end product of the purine catabolic pathway in most mammals (except humans) and can be found in the serum of birds [5,69]. In reptiles, the end product of the purine catabolic pathway is not allantoin, but uric acid [5]. Absence of allantoin in the snake gastrointestinal tract could explain why a substantial number of *S. enterica* clade B were found in snakes, and supports the idea that clade B *Salmonella* are predominantly found in non-allantoin producing hosts [70]. However, we identified one clade B isolate as an exception, isolate 11L-2351 which was sampled from a non-venomous reptile. This isolate belongs to the serovar Montevideo, which is frequently associated with outbreaks of human salmonellosis and shows a broad host range, unlike the other clade B serovars [71–73].

It is possible that the gain and loss of the allantoin catabolism genes may provide an indication of host specificity. The relationship between the pseudogenization of the allantoin metabolic genes and niche adaptation has also been proposed for the invasive non-Typhoidal *Salmonella* (iNTS) reference isolate for *S*. Typhimurium: D23580 [74,75]. Compared with *S*. Typhimurium isolate 4/74, which shows a broad host range, D23580 is unable to utilise allantoin as a carbon source, consistent with the adaptation of invasive *Salmonella* in Africa towards non-allantoin producing hosts [74,75]. Furthermore, the presence of pseudogene accumulation has been described in the allantoin degradation pathway in host-restricted *Salmonella* serovars which cause invasive disease. Thus the loss of ability to grow on allantoin is considered to be a marker of a switch from enteric to invasive disease [69]. These findings may reflect the clinical observation that snake-acquired *Salmonellosis* causes a more invasive disease that commonly results in hospitalisation, compared to non-allantoin producing hosts [20].

## Conclusion

Reptiles are known to harbour a diverse range of *Salmonella* bacteria, however *Salmonella* carriage in a substantial proportion of reptile species has never been investigated. Here we have shown that venomous snakes harbour and shed a medically significant range of *Salmonella* serovars that represents much of the spectrum of the *Salmonella* genus and are phylogenetically distributed in a similar way to *Salmonella* found in non-venomous reptiles. Furthermore, we have identified that venomous snakes epidemiologically-linked to the African continent carry and excrete *Salmonella* serovars which cause human disease in Africa. In one case, the *Salmonella* isolate was resistant to first line antimicrobial agents. It is likely that venomous snakes represent a previously uncharacterised reservoir for *Salmonella* both in captive settings and in the African environment.

Furthermore, we have shown that reptiles are an ideal population of animals for the study of genus-level evolution of *Salmonella*. Reptiles represent a single ecological niche, which carry phylogenetically diverse isolates from the majority of *Salmonella* subspecies. By demonstrating the phenotypic and genotypic conservation of metabolic properties across three phylogenetic groups of *Salmonella* we have shed new light on the evolution of *Salmonella* serotypes.

## Acknowledgements

We are grateful to present and former members of the Hinton lab and Lab H at the Institute of Integrative Biology for helpful discussions. In particular we would like to thank Alex V Predeus and Evelien M Adriaenssens for their ongoing bioinformatic advice and support. We are also grateful to Blanca Perez-Sepulveda for managing the 10 000 *Salmonella* genomes project, without whom sequencing of these isolates would not have been possible. We thank the Centre for Snakebite Research and Interventions at the Liverpool School of Tropical Medicine for access to faecal samples from venomous snakes and to the Leahurst diagnostics unit at the University of Liverpool for providing access to the collection of *Salmonella* isolates from reptiles housed at Chester Zoo.

## Supporting Information Captions

**S1 Table. Reptile-associated-Salmonella strain metadata and control strains metadata.** The table includes all metadata associated with the venomous snake, non-venomous reptile and control strains used throughout this research including details on accessing raw reads were applicable.

**S2 Table. Contextual reference genomes metadata and accession Numbers.** The table lists the strains used in phylogenetic analysis to put the study strains into wider context and includes some basic information in addition to accession numbers.

**S3 Table. QUAST assembly statistics.** The table provides details on assembly statistics for each sequenced genome and highlights those strains which failed quality control in red.

**S4 Table. Carbon source specific requirements.** The table contains a list of the carbon sources used to determine the metabolic capacity of reptile associated *Salmonella*. Specific conditions, concentrations and references are given for each.

**S1 Text. Carbon source utilisation gene sequences.** A full list of the sequences used to determine the presence or absence of genes involved in carbon source utilisation in reptile associated *Salmonella*.

**S1 Figure. The diversity of *Salmonella* isolated from a collection of venomous snakes and non-venomous reptiles including *S. bongori* outgroup.** Core genome maximum likelihood phylogenetic tree. Tree was rooted using *S. bongori* shown here. 25 contextual reference genomes representing previously sequenced isolates from each *Salmonella* subgroup are indicated in purple.

